# Identification of Hub Genes and Functional Annotation of Glaucoma Using Bioinformatic Analysis

**DOI:** 10.1101/2023.10.25.563916

**Authors:** Pavithren Aaron, Navanithan Sivanananthan, Jaspreet Kaur, Jeyvisna Arumainathan, Venkataramanan Swaminathan

## Abstract

Glaucoma is a common eye condition that damages the optic nerve, leading to vision loss and potential blindness if not treated. It affects around 80 million people worldwide and is a major cause of irreversible blindness. This study aims to identify key genes and their functions in glaucoma using bioinformatic analysis. The research follows a five-step approach, including data retrieval, processing, identification of differentially expressed genes (DEGs), protein interaction analysis, functional enrichment analysis, and identification of hub genes. Three GSE datasets (GSE 4316, GSE 53985, and GSE 7144) were selected, resulting in a total of 632 upregulated and 625 downregulated genes. Analysis using the DAVID server revealed significant biological processes related to gene regulation, transcription, and cell adhesion. The protein networks for upregulated and downregulated genes consisted of 581 nodes and 1713 edges, and 575 nodes and 2428 edges, respectively. The Cytohubba plugin identified 10 significant genes for each group. The upregulated genes included PTPRC, INS, SRC, CXCR, FN1, TNF, ACTB, AKT1, ALB, and PECAM1, while the downregulated genes included VEGFA, FN1, TP53, ACTB, AKT1, CCND1, CTNNB1, MYC, CD44, and EGFR. These findings enhance our understanding of glaucoma’s molecular biology and provide valuable insights for prognosis prediction and the development of treatments and medications.

## 1. Introduction

A set of optic neuropathies known as glaucomas are distinguished by the gradual degradation of retinal ganglion cells. These are neurons of the central nervous system, whose axons exit the optic nerve and whose cell bodies are located in the inner retina. These nerves deteriorate, causing cupping, an obvious change in the optic disc’s appearance, and vision loss. [1]

According to Nickells RW, the biological basis of glaucoma is poorly understood and the factors contributing to its progression have not been fully characterised. [2] There are four general categories of adult glaucoma: primary open-angle and angle-closure, and secondary open and angle-closure glaucoma. According to the Centers for Disease Control and Prevention (CDC), open-angle glaucoma is the most common form of glaucoma worldwide. [3]

Globally, glaucoma affects more than 70 million people, 10% of whom are bilaterally blind, and is the leading cause of permanent blindness. There is a significant probability that more people have glaucoma than is currently believed, as the ailment can go untreated until it is advanced [4] According to a study published in The Scientific World Journal, about 50% of the world’s glaucoma cases come from the Asian population, and in Malaysia itself, the prevalence of glaucoma is increasing [5]

Unless early glaucoma symptoms are identified during a regular eye exam, the patient with POAG is frequently asymptomatic until the optic nerve damage is severe. Contrarily, acute angle-closure glaucoma can appear rapidly and cause a more rapid loss of vision along with concomitant headache, nausea, emesis, corneal edema, and eye pain. The most common cause of secondary glaucoma, which results in increased intraocular pressure (IOP) and associated visual neuropathy, is a previous eye injury or disease state. The final variety of glaucoma is normal or low-tension, in which individuals exhibit the same pattern of vision loss as POAG but with normal intraocular pressure readings. [6]

Primary open-angle glaucoma (POAG) is the most common form of glaucoma. When it comes to treatment, the primary goal is to lower intraocular pressure (IOP) to prevent further damage to the optic nerve. Eye drops are commonly prescribed as the first line of treatment for POAG. These medications work by either reducing the production of aqueous humor or improving its outflow to decrease the pressure in the eye. In cases where eye drops alone are not sufficient, other treatment options such as laser trabeculoplasty or surgery may be recommended to enhance the drainage of fluid from the eye. Surgical procedures like trabeculectomy or minimally invasive glaucoma surgery (MIGS) aim to create a new drainage channel or improve the existing one.

The biological mechanism underlying POAG involves a gradual blockage or dysfunction of the trabecular meshwork, the primary drainage pathway for aqueous humor. As the meshwork becomes compromised, the outflow of fluid is impeded, leading to a buildup of pressure within the eye. This increased pressure gradually damages the optic nerve, resulting in vision loss if left untreated. By targeting the reduction of IOP, treatment for POAG aims to slow down or halt the progression of the disease and preserve vision.

According to a journal by Janey Wiggs, glaucoma can be inherited as a complex multifactorial trait, a mendelian autosomal-dominant or autosomal-recessive trait, or both. Some Mendelian forms of the disease have been defined by genetic approaches, which have also helped pinpoint the chromosome locations of genes that are likely to play a role in common complex forms. Future research should focus on finding new glaucoma genes, identifying the clinical phenotypes connected to particular genes and mutations, looking into potential environmental triggers, looking into gene-environment and gene-gene interactions, and creating a database of mutations that can be used for prognostic and diagnostic testing. [7] Therefore the work of this paper is to attempt to identify those said genes that are linked to glaucoma using bioinformatic analysis in hopes to find key information behind the mysteries of the aetiology and potentially making a cure.

To elaborate more on the purpose of this research, bioinformatic analysis has overtime transformed to become an important tool for identifying genes and pathways involved in complex yet persistent diseases such as glaucoma because it allows researchers to analyse large amounts of data from various sources.The analysis can help us identify key genes and pathways that are involved in the disease process and can provide insights into potential therapeutic targets. In addition, bioinformatic analysis can help researchers identify novel biomarkers for diagnosis and prognosis of complex diseases. By using bioinformatic analysis, future researchers can integrate data from multiple sources such as genomics, transcriptomics, proteomics, and metabolomics to gain a more comprehensive understanding of complex diseases.

## 2. Materials and Methods

### Retrieval of data from NCBI

NCBI (https://www.ncbi.nlm.nih.gov/) is a freely accessible website which provides access to biomedical and genomic information. The data of glaucoma was extracted from NCBI under the GEO Datasets database. The results were filtered to expression profiling by array and specified for homo sapiens. GSE 4316, GSE 53985 and GSE 7144 were chosen from the search results. The GSE 4316 has 8 samples, GSE 53985 contains 6 samples while GSE 7144 has 6 samples as well.

### Data Processing and Differentially Expressed Genes ( DEGs ) Selection

The gene expression of the data extracted from NCBI is conducted separately by the GEO2R tool. The GEO2R tool incorporates Bioconductor/R, GEOquery and Limma packages which are used to analyse the differential GEO data expression **[14]**. The DEGs are screened based on the standard value p<0.05 and logFC>1. The upregulated and downregulated DEGs are divided according to logFC positive and logFC negative values.

### Functional Enrichment Analysis

The Database for Annotation, Visualization and Integrated Discovery ( DAVID ) ( https://david.ncifcrf.gov/ ) is a comprehensive set of functional annotation tools which are beneficial to do functional annotation and visualise genes on BioCarta and KEGG pathway maps.**[4]** Once the repeating upregulated and downregulated genes of both GSE IDs are removed, they are submitted in the DAVID server to obtain the functional annotation, Gene Ontology and KEGG pathway of the genes.

### Protein to Protein Interaction Analysis using STRING

STRING is an online database ( https://string-db.org/ ) used to compare DEGs based on the dataset’s results. It provides a comprehensive PPI network. Cut off >0.4 was set as a high confidence interaction score to avoid error in the production of protein networks. Protein to protein interaction network for both upregulated and downregulated genes were obtained.

### Hub genes identification in Cytoscape v 3.9.1

Cytoscape software v 3.9.1( https://cytoscape.org/ ) is a software platform used to visualise complex networks and integrate these with any types of data attribute. Cytohubba app was installed in Cytoscape using the App Manager. The 10 hub genes were generated for upregulated and downregulated genes respectively using Cytohubba in Cytoscape by setting the maximum interactions to 10 and confidence cutoff score to 0.4. The network was extracted in the Maximal Clique Centrality ( MCC ) method which has the highest performance for protein to protein network. The top 10 hub genes are the potential biomarkers for glaucoma.

## 3. Results and Discussion

### Data Collection and Identification of DEGs

From 35 search results, we had selected three studies of similar stature which are GSE 4316, GSE 53985 and GSE 7144 datasets that contained study profiles for gene expression analysis of glaucoma based on source name and cell line.

The Log2FC>1.0 and p-values 0.05 cut-off values were used as the standard primary values to identify the DEGs in the GEO2R tool. The top 250 DEGs were found using the GEOquery.

The identifying DEGs for upregulated and downregulated genes were calculated using logFC 1.0 and logFC 1.0. Between the selected GEO datasets, there were a total of 632 for upregulated and 625 for downregulated genes. The GSE 4316 had a total of 8 samples of glaucoma, GSE 53985 had a total of 6 samples of glaucoma and GSE 7144 had a total of 6 samples of glaucoma. Table 1 lists the details of the datasets that were gathered.

**Table 1:** Lists the details of the datasets that were gathered.

| Sample ID | Total sample | Disease | Platform |
| --- | --- | --- | --- |
| GSE 4316<br>(*PAGE 2 no 25*) | 8 | POAG Glaucoma | GPL570 [HG-U133_Plus_2]<br>Affymetrix Human Genome U133 Plus 2.0 Array |
| GSE 53985<br>(*PAGE 1 NO.7*) | 6 | POAG Glaucoma | GPL570 [HG-U133_Plus_2]<br>Affymetrix Human Genome U133 Plus 2.0 Array |
| GSE 7144<br>(*PAGE 2 no22*) | 6 | POAG Glaucoma | GPL96 [HG-U133A]<br>Affymetrix Human Genome U133A Array |

### Functional Enrichment Analysis

Database for Annotation, Visualization and Integrated Discovery (DAVID’s) functional analysis was obtained to extract the DEGs. The approximation terms for biological process (BP), molecular function (MF), and cellular components (CC) that were upregulated and downregulated were improved by the Gene Ontology (GO) keywords and pathways. A GO biological process was produced using the dataset. The modified Fisher precise EASE score of 0.05 with a p-value of 0.05 and an FDR value of 0.05 was used to determine a strong enrichment value.

The GO biological process of upregulated enriched signal transduction, cell adhesion and RNA polymerase II promoter which is responsible for both positive and negative transcriptional regulation. Upregulated genes achieved protein binding, identical protein binding, DNA binding and protein homodimerization activity for the molecular function of GO. Plasma membrane, integral component of membrane, cytoplasm and cytosol were represented as cellular components.

The DEGs of biological process downregulated included positive regulation of transcription from RNA polymerase II promoter, cell adhesion, DNA-templated and negative regulation of apoptotic process. Besides, the molecular function of GO gained protein binding, identical protein binding, DNA binding, and calcium ion binding. Then, the GO cellular component displayed the nucleus, plasma membrane, cytosol and cytoplasm. These are shown in Table 2.

**Table 2:** Gene Ontology (GO) terms such as biological process (BP), molecular functions (MF), and cellular component (CC) of DEGs by using DAVID.

| Expression | Category | Term | Count | % | P-value | FDR |
| --- | --- | --- | --- | --- | --- | --- |
| **Upregulated** | GOTERM_BP_DIRECT | Signal transduction | 67 | 11.2 | 7.5E-7 | 2.2E-3 |
|  | GOTERM_BP_DIRECT | Positive regulation of transcription from RNA polymerase II promoter | 49 | 8.2 | 9.6E-3 | 8.3E-1 |
|  | GOTERM_BP_DIRECT | Negative regulation of transcription from RNA polymerase II promoter | 36 | 6.0 | 9.2E-2 | 1.0E0 |
|  | GOTERM_BP_DIRECT | Cell adhesion | 31 | 5.2 | 5.2E-4 | 2.6E-1 |
|  | GOTERM_MF_DIRECT | Protein binding | 407 | 68.1 | 1.3E-4 | 6.8E-2 |
|  | GOTERM_MF_DIRECT | Identical protein binding | 60 | 10.0 | 8.3E-2 | 1.0E0 |
|  | GOTERM_MF_DIRECT | DNA binding | 49 | 8.2 | 6.6E-2 | 1.0E0 |
|  | GOTERM_MF_DIRECT | Protein homodimerization activity | 38 | 6.4 | 7.2E-4 | 1.2E-1 |
|  | GOTERM_CC_DIRECT | Plasma membrane | 210 | 35.1 | 4.4E-9 | 2.2E-6 |
|  | GOTERM_CC_DIRECT | Integral component of membrane | 183 | 30.6 | 9.7E-4 | 4.1E-2 |
|  | GOTERM_CC_DIRECT | cytoplasm | 173 | 28.9 | 6.3E-2 | 5.9E-1 |
|  | GOTERM_CC_DIRECT | Cytosol | 170 | 28.4 | 6.2E-2 | 5.9E-1 |
| **Downregulated** | GOTERM_BP_DIRECT | Positive regulation of transcription from RNA polymerase II promoter | 57 | 9.6 | 1.3E-4 | 2.8E-2 |
|  | GOTERM_BP_DIRECT | Cell adhesion | 46 | 7,8 | 1.7E-10 | 1.8E-7 |
|  | GOTERM_BP_DIRECT | Positive regulation of transcription, DNA-templated | 39 | 6.6 | 1.6E-4 | 3.1E-2 |
|  | GOTERM_BP_DIRECT | Negative regulation of apoptotic process | 34 | 5.7 | 1.2E-5 | 3.7E-3 |
|  | GOTERM_MF_DIRECT | Protein binding | 415 | 70.1 | 8.7E-6 | 1.1E-3 |
|  | GOTERM_MF_DIRECT | Identical protein binding | 65 | 11.0 | 1.9E-2 | 5.1E-1 |
|  | GOTERM_MF_DIRECT | DNA binding | 49 | 8.3 | 6.9E-2 | 1.0E0 |
|  | GOTERM_MF_DIRECT | Calcium ion binding | 42 | 7.1 | 9.2E-5 | 7.1E-3 |
|  | GOTERM_CC_DIRECT | Nucleus | 193 | 32.6 | 2.8E-3 | 6.6E-2 |
|  | GOTERM_CC_DIRECT | Plasma membrane | 177 | 29.9 | 1.2E-3 | 3.8E-2 |
|  | GOTERM_CC_DIRECT | Cytosol | 177 | 29.9 | 5.8E-3 | 1.0E-1 |
|  | GOTERM_CC_DIRECT | Cytoplasm | 169 | 28.5 | 7.0E-2 | 6.2E-1 |

The KEGG pathway of upregulated DEGs gained ras signaling pathway, neutrophil extracellular trap formation, vascular smooth muscle contraction and systemic lupus erythematosus. The downregulated pathway is PI3K-Akt signaling pathway, pathways in cancer, human papillomavirus infection and proteoglycans in cancer. Using DAVID v6.8 online tools, Table 3 displays the KEGG pathway analysis of differentially expressed genes linked to glaucoma.

**Table 3:** KEGG pathway analysis of differentially expressed genes associated with Glaucoma by using DAVID online tools.

| Expression | Term | Count | % | P-value | FDR |
| --- | --- | --- | --- | --- | --- |
| **Upregulated** | Ras signaling pathway | 19 | 3.2 | 1.5E-3 | 6.2E-2 |
|  | Neutrophil extracellular trap formation | 17 | 2.8 | 9.5E-4 | 5.6E-2 |
|  | Vascular smooth muscle contraction | 16 | 2.7 | 6.2E-5 | 1.1E-2 |
|  | Systemic lupus erythematosus | 16 | 2.7 | 7.3E-5 | 1.1E-2 |
| **Downregulated** | Pathways in cancer | 41 | 6.9 | 2.9E-7 | 1.2E-5 |
|  | PI3K-Akt signaling pathway | 30 | 5.1 | 2.8E-6 | 1.0E-4 |
|  | Human papillomavirus infection | 28 | 4.7 | 6.8E-6 | 2.0E-4 |
|  | Proteoglycans in cancer | 26 | 4.4 | 7.2E-9 | 1.1E-6 |

### Protein to Protein Interaction Analysis using STRING

The PPI of DEGS downregulated and upregulated were evaluated using the STRING tool as shown in Figure 1 and Figure 2. The protein network of the upregulated gene set has 581 nodes and 1713 edges while the down regulated protein network has 575 nodes and 2428 edges.

**Figure 1:**
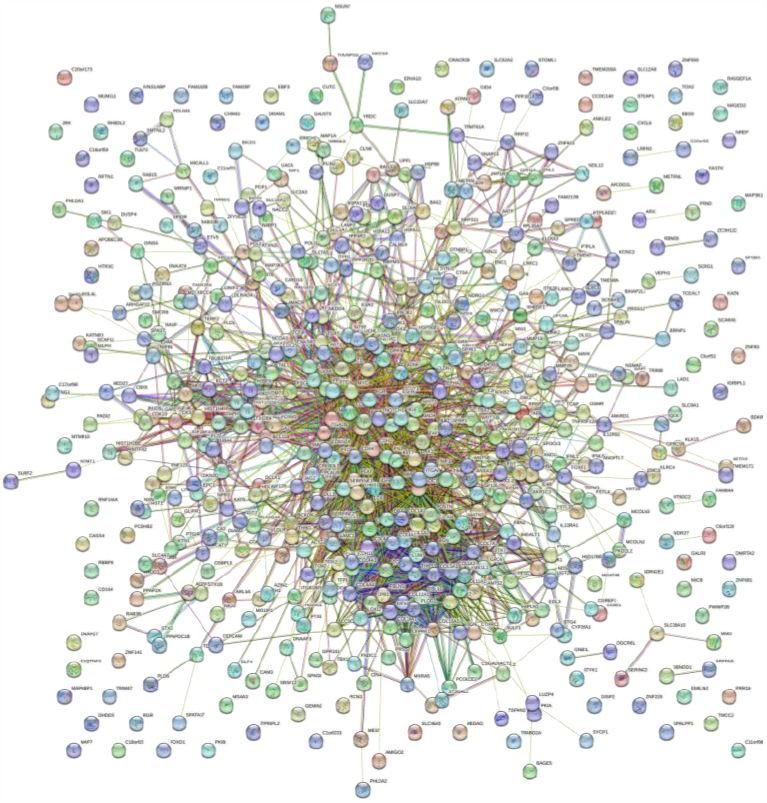
The picture above shows the protein to protein interaction in a downregulated gene set in STRING.

**Figure 2:**
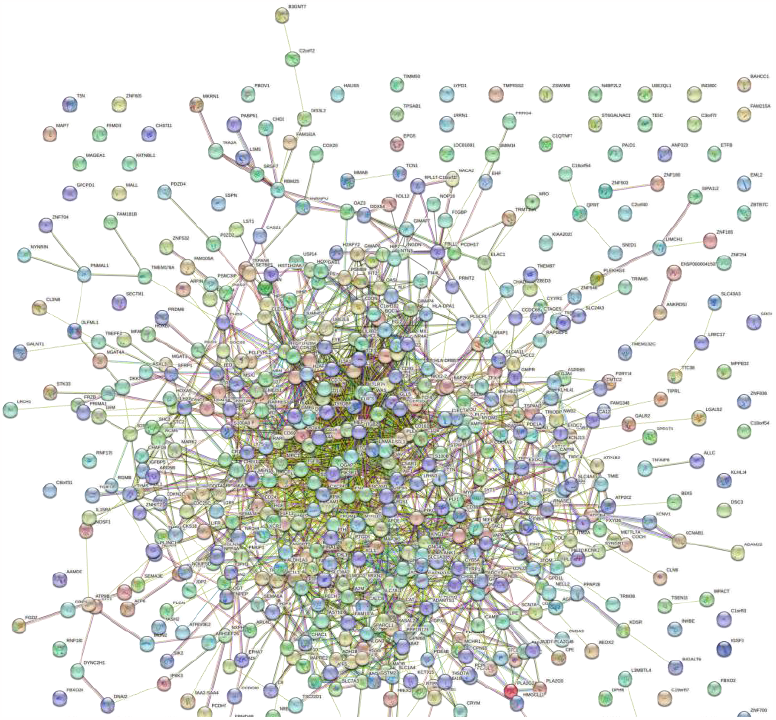
The picture above shows the protein to protein interaction between the upregulated gene set in STRING.

### Top 10 Hub Genes Identification using Cytoscape

The top 10 hub genes of Glaucoma are identified by using Cytohubba, plugged in the Cytoscape. The top 10 hub genes for upregulated gene sets are shown in Figure 3 while the top 10 hub genes for downregulated gene sets are shown in Figure 4. The results show the display names of the hub genes.

**Figure 3:**
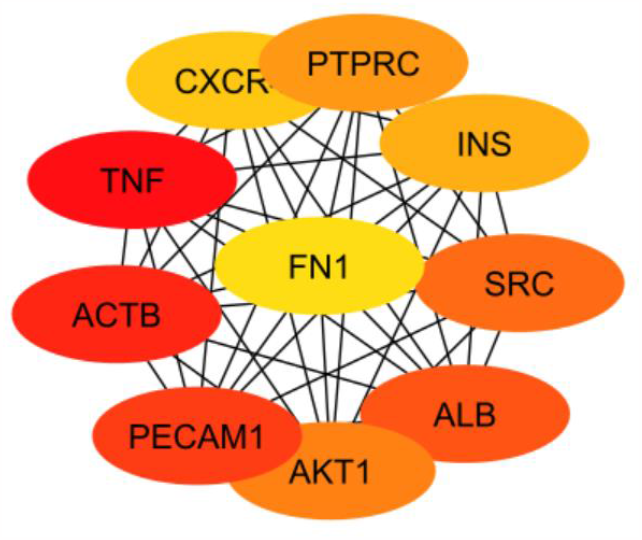
The picture above shows the top 10 hub genes of upregulated Glaucoma gene set obtained from Cytoscape.

**Figure 4:**
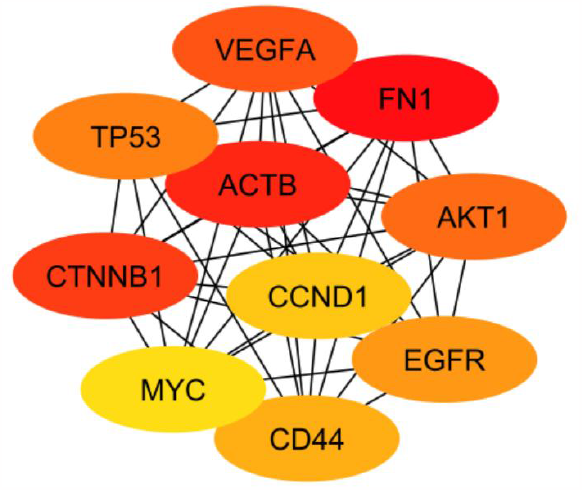
The picture above shows the top 10 hub genes of downregulated Glaucoma gene set obtained from Cytoscape.

Based on the results, GSE 4316, GSE 53985 and GSE 7144 was the data that was collected on glaucoma. The selected GEO datasets contained 625 downregulated genes and 632 upregulated genes in total.

In this study, DEGs were found in the glaucoma biology process (BP), cellular component (CC), and molecular function (MF) in order to obtain functional annotation. This was made possible by Gene Ontology (GO) terms upregulate and downregulate. Based on the Database for Annotation, Visualization and Integrated Discovery (DAVID) server, it has been shown that there is biological process in the positive and negative regulation of transcription from RNA polymerase II promoter, regulation of transcription, DNA-templated and cell adhesion. As for the cellular component, it is shown that the gene could be detected in the nucleus, cytosol, cytoplasm and plasma membrane. It has also shown that in the molecular function, it shows the identical protein binding, protein binding, DNA binding and calcium ion binding.

The PPI network helps to identify the interactions between proteins and its role in glaucoma. Based on the figure 1, the protein network of the upregulated gene set has 581 nodes and 1713 edges while the down regulated protein network has 575 nodes and 2428 edges. Moreover, the top 10 hub genes responsible for the development of Glaucoma are identified using Cytoscape for both upregulated and downregulated genes. The hub genes for upregulated genes are PTPRC, INS, SRC, CXCR, FN1, TNF, ACTB, ATK1, ALB and PECAM1. As for the downregulated hub genes that were obtained are VEGFA, FN1, TP53, ACTB, AKT1, CCND1, CTNNB1, MYC, CD44 and EGFR. All the above genes have its functions causing Glaucoma. For instance, PTPRC is also known as Protein Tyrosine Phosphatase, Receptor Type, C. It is a signalling molecule which can regulate cellular processes. It is an essential regulator for T - cell and B - cell signalling. Moreover, INS, known as insulin, is essential for the regulation of carbohydrate and lipid metabolism.

TNF is also another hub gene of upregulated genes. The gene encodes a multifunctional proinflammatory cytokine that belongs to the tumour necrosis factor (TNF) superfamily. This cytokine is involved in the regulation of a wide spectrum of biological processes including cell proliferation, differentiation, apoptosis, lipid metabolism, and coagulation. In relation to cancer biology, TNF was initially reported as a serum factor that induced necrosis of tumours **[3]**.

Actin Beta is a protein coding gene. This gene encodes one of six different actin proteins. Actins are highly conserved proteins that are involved in cell motility. Actin is a highly conserved protein that polymerizes to produce filaments that form a cross-linked network in the cytoplasm of cells **[9]**. Moreover, AKT serine/threonine Kinase 1 gene encodes one of the three members of the human AKT serine-threonine protein kinase family which are often referred to as protein kinase B alpha, beta and gamma. AKT/PI3K forms a key component of many signalling pathways that involve the binding of membrane-bound ligands such as receptor tyrosine kinases, G-protein coupled receptors and integrin-linked kinases. AKT1 is one of 3 closely related serine/threonine-protein kinases (AKT1, AKT2 and AKT3) called the AKT kinase and which many processes including metabolism, proliferation, cell survival, growth and angiogenesis **[10]**.

VEGFA is a member of the PDGF/VEGF growth factor family. It encodes a heparin-binding protein, which exists as a disulfide-linked homodimer. This gene is upregulated in many known tumours and its expression is correlated with tumour stage and progression. Activated VEGFA signalling pathways can promote proliferation and migration of endothelial cells as well as their survival and vascular permeability **[11]**.

A Tumour Protein 53, TP53, is a protein coding gene. This gene encodes a tumour suppressor protein containing transcriptional activation, DNA binding and Oligomerization domains. The encoded protein responds to diverse cellular stresses to regulate expression of target genes, thereby inducing cell cycle arrest, apoptosis, senescence, DNA repair or changes in metabolism. TP53 mutation might adversely affect the survival of patients treated with targeted therapy for oncogenic driver mutations in lung cancer. **[12]**

Catenin Beta 1 protein encoded by this gene is part of a complex of proteins that constitute adherens junctions (AJs). Beta-catenin protein is an integral part of the canonical Wnt signalling pathway. In the presence of Wnt ligand, CTNNB1 is not ubiquitinated and accumulate in the nucleus, where it acts as coactivator for transcription factors of the TCG/LEF family, leading to activate Wnt responsive genes **[11]**.

MYC is a proto-oncogene and encodes a nuclear phosphoprotein that plays a role in cell cycle progression, apoptosis and cellular transformation. The encoded protein forms a heterodimer with related transcription factor MAX. Activates the transcription of growth-related genes **[13]**. EGFR or known as Epidermal Growth Factor Receptor is widely recognized for its importance in cancer. This protein encoded by this gene is transmembrane glycoprotein that is a member of the protein kinase superfamily. This protein is a receptor for members of the epidermal growth factor family.

Albumin is a gene which encodes for protein in human blood. The protein regulates blood plasma colloid osmotic pressure. The diseases associated with Albumin include Analbuminemia and Hyperthyroxinemia. Furthermore, fibronectin is a glycoprotein in soluble dimeric form in plasma. It can cause diseases such as Spondylometaphyseal Dysplasia, Corner Fracture Type and Glomerulopathy.

## 4. Conclusions

In conclusion, 632 upregulated and 625 downregulated genes were among the 750 DEGs in the glaucoma gene expression profiling datasets GSE 4316, GSE 53985 and GSE 7144. These were incorporated using the bioinformatics technique. STRING and CytoScape used CytoHubba to incorporate the top 10 glaucoma hub genes into the PPI network. The top 10 significant genes that were upregulated were PTPRC, INS, SRC, CXCR, FN1, TNF, ACTB, AKT1, ALB, PECAM1, while the top 10 significant genes that were downregulated were VEGFA, FN1, TP53, ACTB, AKT1, CCND1, CTNNB1, MYC, CD44, EGFR. These support a deeper understanding of the molecular biology of biomarkers hub genes in glaucoma and provide precise techniques for prognosis prediction. Understanding the mechanism of glaucoma in greater details was useful for determining risk, developing medications and finding treatments.

## Acknowledgments

We, the authors of the paper, would like to show our gratitude towards our affiliation, Management and Science University for providing us with the knowledge and resources to carry out this paper.

